# Direct cell reprogramming by a designed agonist inducing HER2-FGFR proximity

**DOI:** 10.1101/2025.10.12.681903

**Authors:** Riya Keshri, Marc Exposit, Mohamad Abedi, Derrick R Hicks, Zachary Foreman, Ashish Phal, Yen Chian Lim, Philip Barrett, Catherine Sniezek, Jinlong Lin, Thomas Schlichthaerle, Alexander J Robinson, Damien Detraux, Tung Chan Ching, Keija Wu, Brian Coventry, Lemuel Chang, Alec S.T. Smith, David L Mack, Devin K Schweppe, Beatriz Estrada Martin, Kalina Hristova, Julie Mathieu, David Baker, Hannele Ruohola-Baker

## Abstract

Growth factor induced receptor dimerization and activation of downstream pathways can modulate cell fate decisions. Here, we investigate the potential of de novo designed synthetic ligands, termed Novokines, to reprogram cell identity by inducing proximity of novel pairs of receptor subunits. We find that a design, H2F, that brings together HER2 (which has no known natural ligand) and the FGF receptor has potent signaling activity. H2F induces robust signaling and reprograms fibroblasts into myogenic cells. Unlike native FGF ligands, H2F selectively activates the MAPK pathway without engaging PLCγ-mediated Ca²⁺ signaling. FRET assays confirm H2F-mediated HER2–FGFR proximity, and phosphoproteomic analysis reveals activation of MAPK effectors. H2F-induced ERK phosphorylation is abolished in cells expressing a kinase-dead FGFR1 (K514M) mutant, confirming the requirement for FGFR catalytic activity. H2F treatment significantly increases myofiber formation from adult patient–derived primary myoblasts, demonstrating its capacity to promote myogenic regeneration. Our findings demonstrate that synthetic receptor pairings can rewire signaling outputs to drive regeneration, providing a programmable platform for cell fate engineering.

## Introduction

The direct conversion, or transdifferentiation, of somatic non-muscle cells into skeletal myocytes offers substantial therapeutic potential in the future for treating conditions such as muscle atrophy and sarcopenia. Cell fate transitions are often orchestrated by growth factors that induce the assembly of specific receptor pairs, leading to the transactivation of intracellular kinase domains and the initiation of lineage-specific signaling cascades. However, natural growth factors have not been shown to induce fibroblast-to-myoblast conversion, suggesting that new strategies are needed to unlock alternative cell identities.

Advances in de novo protein design have enabled the generation of compact (50–110 amino acids), high-affinity receptor-binding domains with remarkable isoform specificity, thermal stability, and modular architecture. These domains typically act as antagonists in monomeric form but can function as potent agonists when presented in multivalent configurations that promote receptor clustering ^1,2,3^. We reasoned that the unique modularity and precision of these synthetic ligands could be leveraged to reprogram cell fate. We investigated whether bringing together receptor subunits that are not known to naturally dimerize, such as HER2 with an unrelated Receptor Tyrosine Kinase (RTK), could trigger noncanonical signaling events capable of driving lineage conversion.

In this study, utilizing the skeletal muscle transdifferentiation model we identified the HER2:FGFR Novokine (H2F) as a potent driver of myogenesis. H2F promotes HER2-FGFR heterodimerization, confirmed by FRET. H2F signaling relies on FGFR kinase activity and Y653/Y654 phosphorylation, with phosphoproteomics revealing activation of MAPK effectors and regulators such as GAPs and GEFs. H2F stands as an exceptional ligand that can bifurcates FGFR pathways where MAPK/AKT activation and eliminates PLCγ/Ca²⁺ signaling. Functionally, H2F enhances direct reprogramming of fibroblast into skeletal muscle cells, increases myotube formation from patient-derived myoblasts, and also supports iPSC-derived myogenesis. Like FGF2, H2F preserves iPSC pluripotency, indicating PLCγ signaling is dispensable for stemness. But unlike FGF2, H2F does not support endothelial differentiation, highlighting the relevance of Ca²⁺ signaling arm in endothelial cell fate determination. H2F activity could be exploited in various physiological niches where HER2 and FGFR1/2c are co-expressed. Broadly, our modular strategy of fusing designed binders to recruit unrelated endogenous receptor pairs establishes Novokines as reprogramming tools via rewiring novel signaling outputs. Novokines represent a powerful framework for both therapeutic development in regenerative medicine and discovery of fundamentals of signalling logics through receptor remodelling.

## Results

We screened a set of de novo designed Novokines constructed by fusing two computationally designed receptor-binding domains via a flexible linker (Fig. 1A–C; Abedi et al., submitted) for effects on skeletal muscle reprogramming. These binding domains are engineered to target a diverse set of Receptor Tyrosine Kinases and Receptor Serine/Threonine Kinases (RTKs/RSTKs) with high affinity and specificity ^1–3^. By simultaneously engaging two distinct receptor subunits, Novokines bring these receptors into close proximity. This enforced proximity may promote cross-phosphorylation of intracellular domains or associated signaling components, thereby initiating novel downstream signaling events with the potential to influence cell fate reprogramming

**Figure 1:**
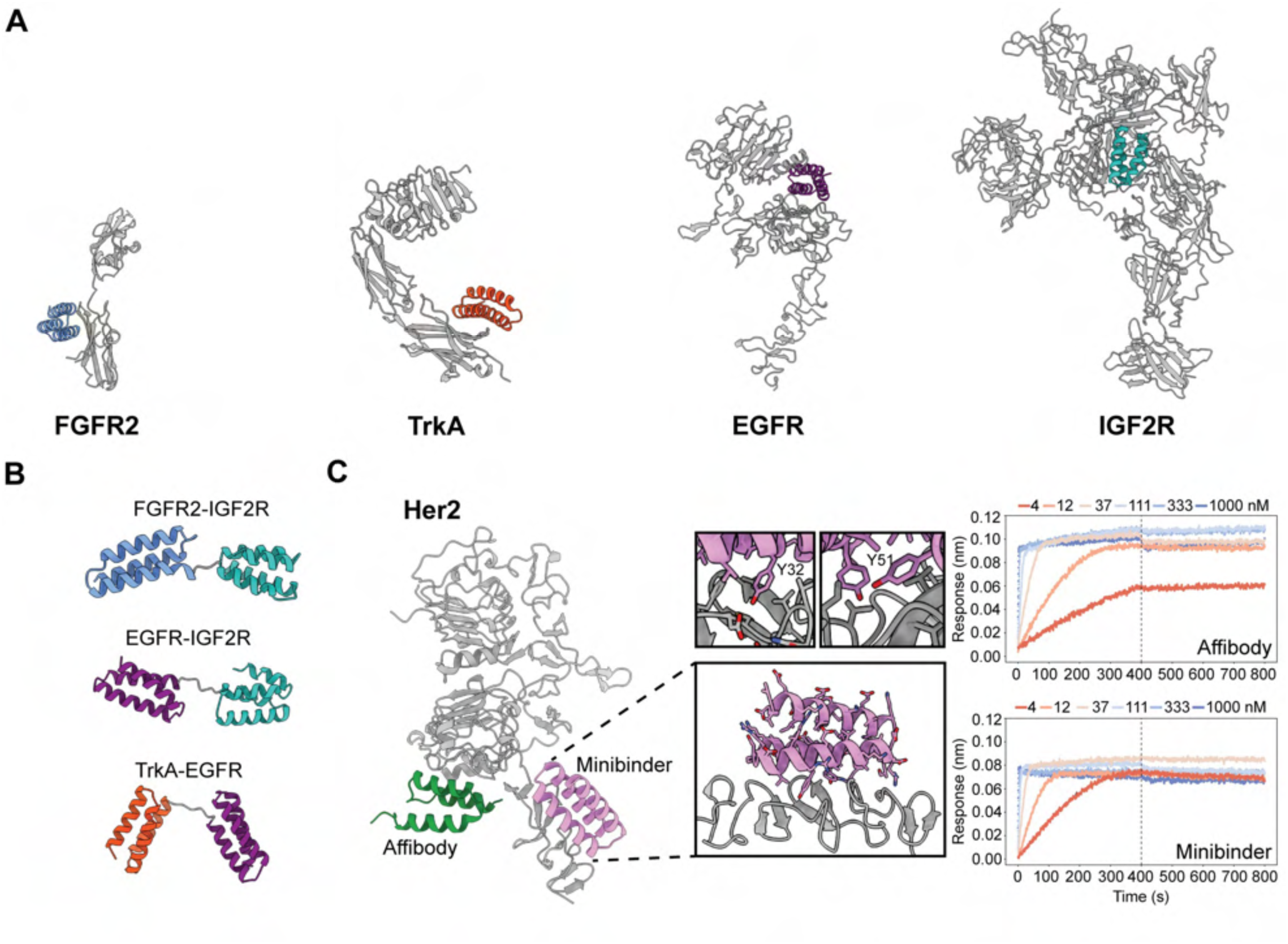
De novo design of novokines. (A) Previously designed minibinders against FGFR2 (PDB ID: 7TYD), TrkA (PDB ID: 7N3T), EGFR (PDB ID: 3NJP) and IGF2R (PDB ID: 6UM2). (B) Flexibly linked novokines. (C) HER2 receptor with Affibody (PDB ID: 3MZW) superimposed together with HER2 minibinder. Zoom ins show key residues important for minibinder-HER2 receptor interaction. Right: Comparison of binding between the affibody (top) and the minibinder (bottom) to the HER2 receptor using BLI measurements.

### Designed Novokines enhanced skeletal muscle reprogramming

We began by developing a screen for compounds that increased the reprogramming efficiency of fibroblasts expressing MyoD. MyoD induces direct reprogramming of human fibroblasts into skeletal muscle ^4,5^, but the efficiency of the conversion process remains low^6,7,8^. Accordingly, even prolonged MyoD overexpression in HFF-iMYOD (a Doxycycline-inducible MyoD overexpression line in human foreskin fibroblasts, Fig. 2A) for 7 or 14 days did not show improvement in full or partial commitment to myogenic fate with the majority of cells exiting the reprogramming pathway (Fig. 2B-C). We optimized the assay for screening the library of Novokines (Fig. 2). The expression of several muscle markers, including *MYH3, MYH7, MYH8, TTN3, ACTN2, DESMIN, TITIN, ACTA1* (skeletal muscle-specific actin), shows that fibroblast fate has partially changed towards trans differentiated myotubes (tSKM) (Fig. 2E). We tested natural ligands (FGF2, FGF7, FGF10, IL-6, EGF, BMP4, VEGF, TNF-a) but did not observe significantly enhanced myogenic conversion (Fig. 3D), highlighting the need for synthetic approaches. We therefore screened 155 designed novokines, targeting various combinations of RTKs/RSTKs, added on days 4-8 post-MyoD induction (Fig. 2D), to identify any that enhance myogenic efficiency.

**Figure 2.**
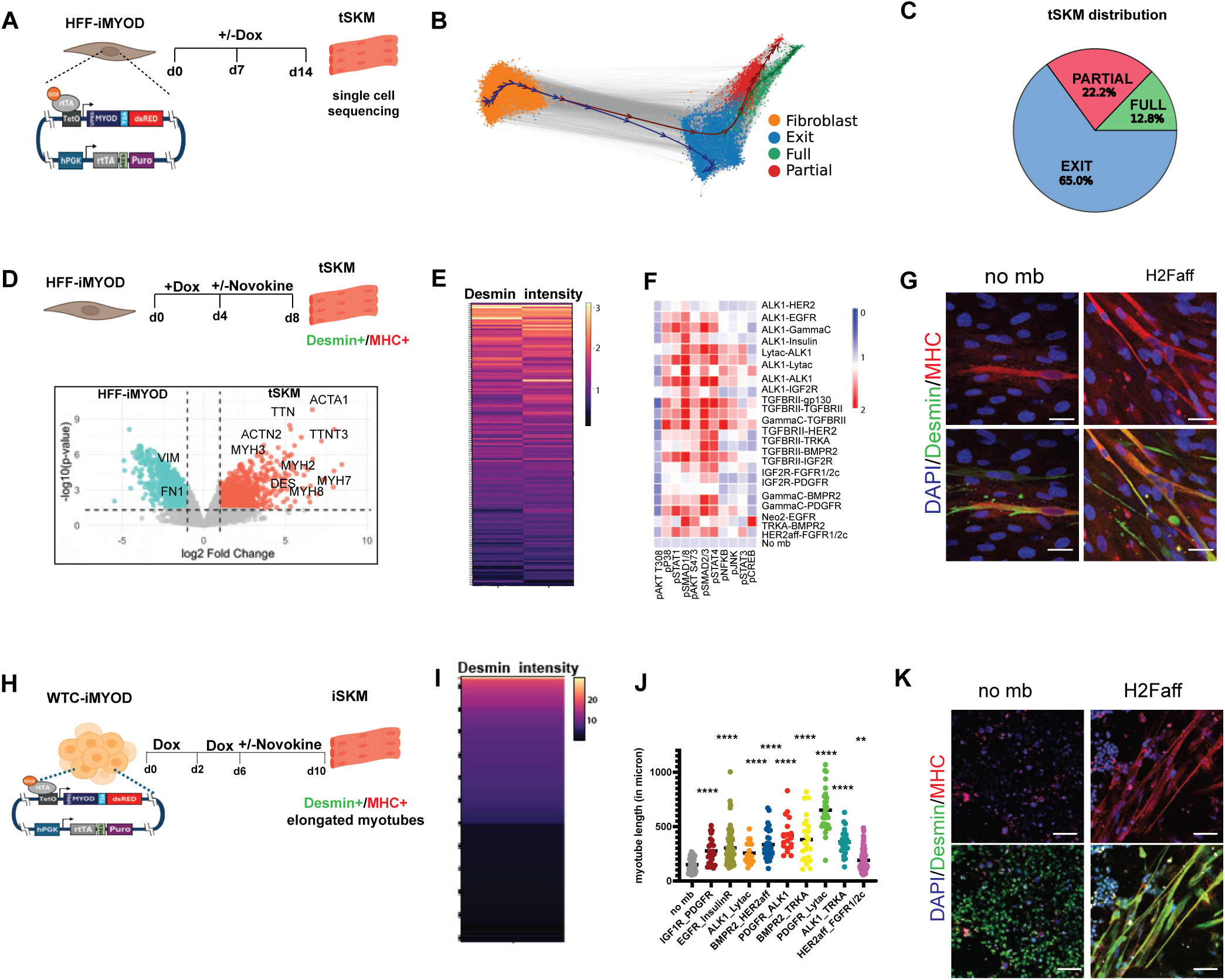
Designed novokines enhance skeletal muscle reprogramming. (A) Schematic showing human foreskin fibroblast (HFF-iMYOD) harvested on d7 and d14 of continuous doxycycline treatment for single cell sequencing. (B) percentage distribution of cells in full myogenic, partial myogenic or exit state post d7 or d14 of MYOD inducible expression. (C) UMAP showing pseudotime trajectory analysis of fibroblast to commitment, partial or exit state post d7 or d14 of MYOD inducible expression. (D) Schematic showing novokine screen method in HFF-iMYOD derived myogenic transdifferentiation. Volcano plot analyzing differential expressed genes in tSKM vs HFF-iMYOD cells in bulk RNA sequencing analysis. (E) Heat map of desmin intensity on immunostaining of tSKMs in novokine treatment in comparison to control (n=2), in two independent primary screens of designed novokines in myogenic transdifferentiation. (F) Heatmap showing various phospho-effectors analysed in flow cytometry of EA.hy926 cells upon novokine treatment for 15 min. (G) Confocal image showing desmin staining (green), MHC (red), and DAPI (blue) in control and HER2mb:FGFRmb (H2F) treated conditions in the myogenic transdifferentiation assay (D). scale bar=30 um (H) Schematic showing of screening method for novokines in human iPSCs-derived myogenic differentiation. (I) Desmin Intensity heat map of iSKMs treated with novokines in comparison to WTC-iMyoD. (J) In various 100nM Novokine treatments that are positive for showing elongated myotube formation are represented. Myotube length has been significantly increased in comparison to no minibinder treatment (measured in micron). (K) Confocal images showing desmin staining(green), MHC(red), and DAPI(blue) in control, H2F treated conditions in the iPSC derived myogenic differentiation assay, scale bar=50 um. ****p < 0.0001

**Figure 3.**
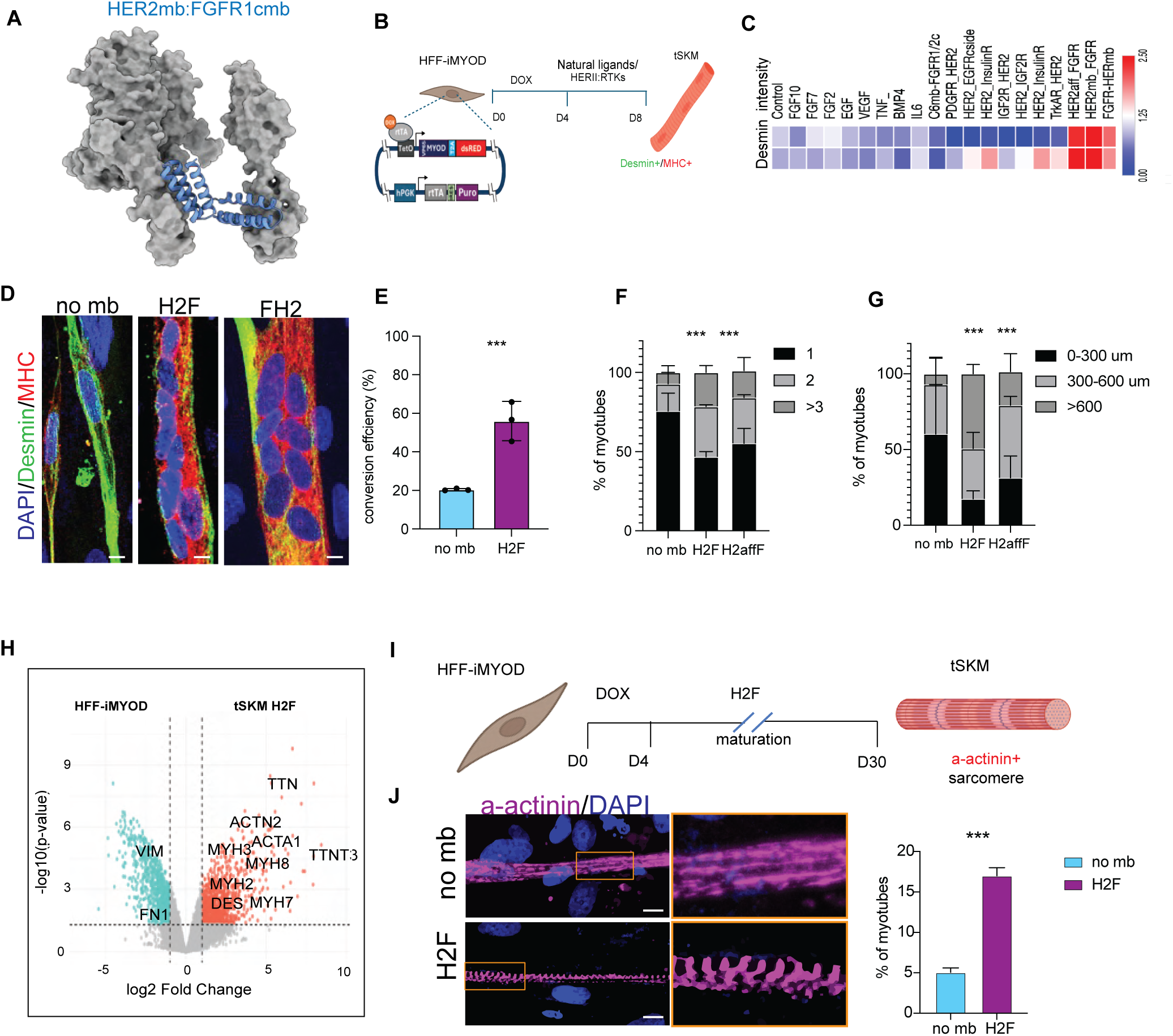
HER2 and FGFR1/2c heterodimerization shows enhanced myogenic reprogramming. (A) AlphaFold-generated predicted structures of the HER2 receptor minibinder fused with a flexible linker to FGFR1c minibinder in the orientation HER2mb:FGFRcmb (blue, H2F) when bound to the extracellular domains of their cognate receptors (grey). (B) Schematic showing screen method for several HER2-RTKs novokines in human foreskin fibroblast (HFF-iMYOD) derived myogenic transdifferentiation. (C) Heatmap showing various natural ligands and HER2aff:RTK novokines, do not increase desmin intensity relative to control whereas H2F, and FGFRmb:HER2mb (FH2mb) shows increased desmin intensity. (D) Confocal images showing desmin staining (green), MHC (red), and DAPI (blue) in control, H2F, and FH2mb treated tSKM. scale bar=5 um (E) Graph showing that H2F treatment increases the conversion efficiency (percentage of nuclei in desmin positive cells) of the myogenic transdifferentiation assay.(F) Graph of the percentage of myotubes with one, two, or three and greater number of nuclei. Shows H2F and H2affF treatment significantly increases multinucleation. (G) Graph showing that H2F/FH2mb/H2affF treated tSKM show increased length compared to control (H) A volcano map showing mRNA expression of various muscle markers are upregulated in tSKM treated with H2F. (I) Schematic showing method for the long-term maturation (30 days) assay of H2F tSKM. (J) Confocal Images of H2F long term maturation tSKM stained for DAPI (blue) and alpha-actinin (magenta), scale bar=5 um. H2F treatment led to significant increase in the maturation of the myotubes as alpha-actinin striation was significantly increased in H2F treated tSKMs. Graph showing percentage of myotubes with alpha-actinin positive sarcomeres. ***p < 0.001

We identified novokines that passed both primary and secondary screens, significantly enhancing myogenic conversion efficiency, as assessed by Desmin and MHC (Myosin Heavy Chain) expression (Fig. 2D, 2G; S2A,2B). To further investigate their mechanisms, we analyzed selected candidate novokines via flow cytometry-based phospho-effectors (pJNK, pP38, pAKT, etc) analysis (Fig. 2E). In starved immortalized EA.hy926 cells (an immortalized human umbilical vein endothelial cell line), a brief 15-minute novokine treatment was sufficient to elicit phospho-effector signaling responses (Fig. 2F). We further identified a subset of novokine treated tSKM which showed increased mitochondrial respiration measured using the seahorse assay (e.g.FGFR_EGFRc, EGFRc_PDGFR, TGFBR2_BMPR2 etc.) without significant change in myogenic conversion efficiency (Fig.S2C). The profound metabolic rewiring induced by these novokines suggests a potential to restore mitochondrial health, offering future opportunities to address mitochondrial-related disorders.

We next evaluated the novokine library in an independent myogenic model using induced pluripotent stem cell (iPSC)-derived skeletal muscle differentiation. To this end, we generated a WTC-11 iMYOD cell line, in which MyoD expression is driven by a doxycycline-inducible promoter ^6^(Fig. 2I). To assess the ability of novokines to enhance myogenic differentiation, we established an iPSC-derived skeletal myocyte (iSKM) assay: MyoD was induced for six days to initiate myogenic programming, followed by four additional days of differentiation in the presence of individual Novokines. We screened the Novokine library for effects on differentiation, using desmin and myosin heavy chain (MHC) expression as molecular markers, along with morphological assessments (Fig. 2I). A subset of novokines significantly increased desmin and MHC levels (Fig. 2I, S2D). Among these, 13 novokines also induced hallmark morphological features of myogenic differentiation, including elongated, multinucleated myotubes (Fig. 2J-K S2E).

In this study, we focus on a specific novokine, HER2aff–FGFRmb (H2affF), which promoted myogenic differentiation in both the fibroblast transdifferentiation assay and the iPSC-derived skeletal muscle model. H2F comprises an FGFR-binding minibinder (FGFRmb) fused to a HER2-specific affibody (H2aff)^1^. To independently test whether bringing HER2 and FGFR into proximity is sufficient to trigger signaling, we engineered a new minibinder for HER2. HER2, a member of the EGF receptor family of receptor tyrosine kinases (RTKs), is known to heterodimerize with EGFR upon ligand binding to initiate downstream signaling^9^. We designed a HER2 minibinder (H2mb) that binds to the extracellular domain IV of HER2-distinct from the affibody binding site (Fig. 1C, S1A) using Rifgen (Cao et al), followed by ProteinMPNN and AlphaFold filtering as outlined by Bennett et al ^10^. From the initial pool of ∼13,000 designs, candidates were screened via yeast surface display (YSD) and ranked by binding affinity, as described previously. The top-performing binder was further optimized through site-saturation mutagenesis (SSM), and enriched variants were incorporated into a degenerate codon combinatorial library, which was again screened by YSD ^3^(Fig. S1C). 96 top variants were assessed for soluble expression in *E. coli* and binding affinity via biolayer interferometry (BLI/Octet) (Fig.1C,1SB). A single lead candidate with robust soluble expression and sub-nanomolar binding affinity was selected for fusion with FGFRmb and subsequent functional characterization.

### HER2mb:FGFR1/2cmb mediated receptor heterodimerization augments muscle Reprogramming

To gain molecular insights into HER2 and FGFR1/2c heterodimerization, we expressed three constructs-HER2mb-FGFRmb (H2F), FGFRmb-HER2mb (FH2), and HER2aff-FGFRmb (H2affF) (Fig. 3A) and compared their effects on myogenic transdifferentiation (Fig. 3B-D). All three novokines enhanced transdifferentiation efficiency. In contrast, neither natural ligands (FGF2, FGF7, FGF10, IL-6, EGF, BMP4, VEGF, TNF-a), designed FGFR1/2c agonist C6-79C-mb7 nor HER2aff fused with other RTK minibinders (such as PDGFRmb, EGFRcmb, IGF2Rmb, TRKA Receptor mb and Insulin Receptor mb) increased transdifferentiation efficiency (Fig. 3C, S3A). These results suggest that HER2:FGFR heterodimerization activates a unique signaling pathway that facilitates myogenic fate conversion of fibroblasts (Fig. 3C, D S3A). Direct muscle reprogramming was achieved with HER2 and FGFR1/2c heterodimerization using two independent HER2 binders (HER2mb and HER2Affibody), despite their distinct binding surfaces and locations within the HER2 extracellular domain (Fig. 1C).

H2F treatment significantly enhanced myogenic conversion efficiency to about 56% compared with control (Fig. 3E). H2F-converted cells also exhibited two times greater multinucleation relative to control cells and formed significantly longer myotubes (≈5-fold increase) (Fig. 3F,G). Additionally, myotubes treated with H2F or H2affF showed higher metabolic activity, characterized by a more elaborate mitochondrial network and increased Mito-Orange uptake (Fig. S3B). Bulk mRNA analysis showed increased expression of various muscle markers in H2F treated myotubes (Fig. 3H). Collectively, these findings indicate that H2F not only increases the efficiency of direct reprogramming but also promotes downstream myogenic differentiation. Myofiber maturation was significantly enhanced when the H2F treatment was extended from 4 days to 30 days, as evidenced by significantly increased formation of well-organized Z-bands of sarcomeric α-actinin in the transdifferentiated muscle cells (Fig. 3I,J, S3C).

Collectively, these data indicate that H2F minibinders effectively promote the conversion of fibroblasts into myotubes and their subsequent maturation. Furthermore, these findings suggest that heterodimerization of HER2 and FGFR receptors activates a distinct pro-myogenic signaling program that is not induced by other HER2 receptor tyrosine kinase pairings or by stimulation of FGFR or HER2 alone.

### HER2mb-FGFRmb induced HER2-FGFR pair drives downstream signaling

We performed quantitative imaging Förster Resonance Energy Transfer (QI-FRET) experiments to assess the interaction between HER2 and FGFR1 in the presence of the synthetic ligand H2F[30]. Experiments were performed with truncated receptors, with the intracellular domains substituted with fluorescent proteins (Fig. 4SA,B). HER2-mEYFP (donor) and FGFR-mCherry (acceptor) were co-expressed in individual CHO cells, and their expression levels were comparable in both H2F-treated and untreated conditions (Fig. 4SB). In the presence of H2F, FRET efficiencies were elevated relative to the no-ligand control (Fig. 4SA; in the latter case FRET efficiencies were still higher than baseline [31], suggesting possible interactions between HER2 and FGFR1 in the absence of ligand. Comparison of the deviation from “proximity FRET (Fig. 4A), shows a statistically-significant increase due to H2F, indicating that H2F stabilizes the HER2–FGFR heterocomplex .

**Figure 4.**
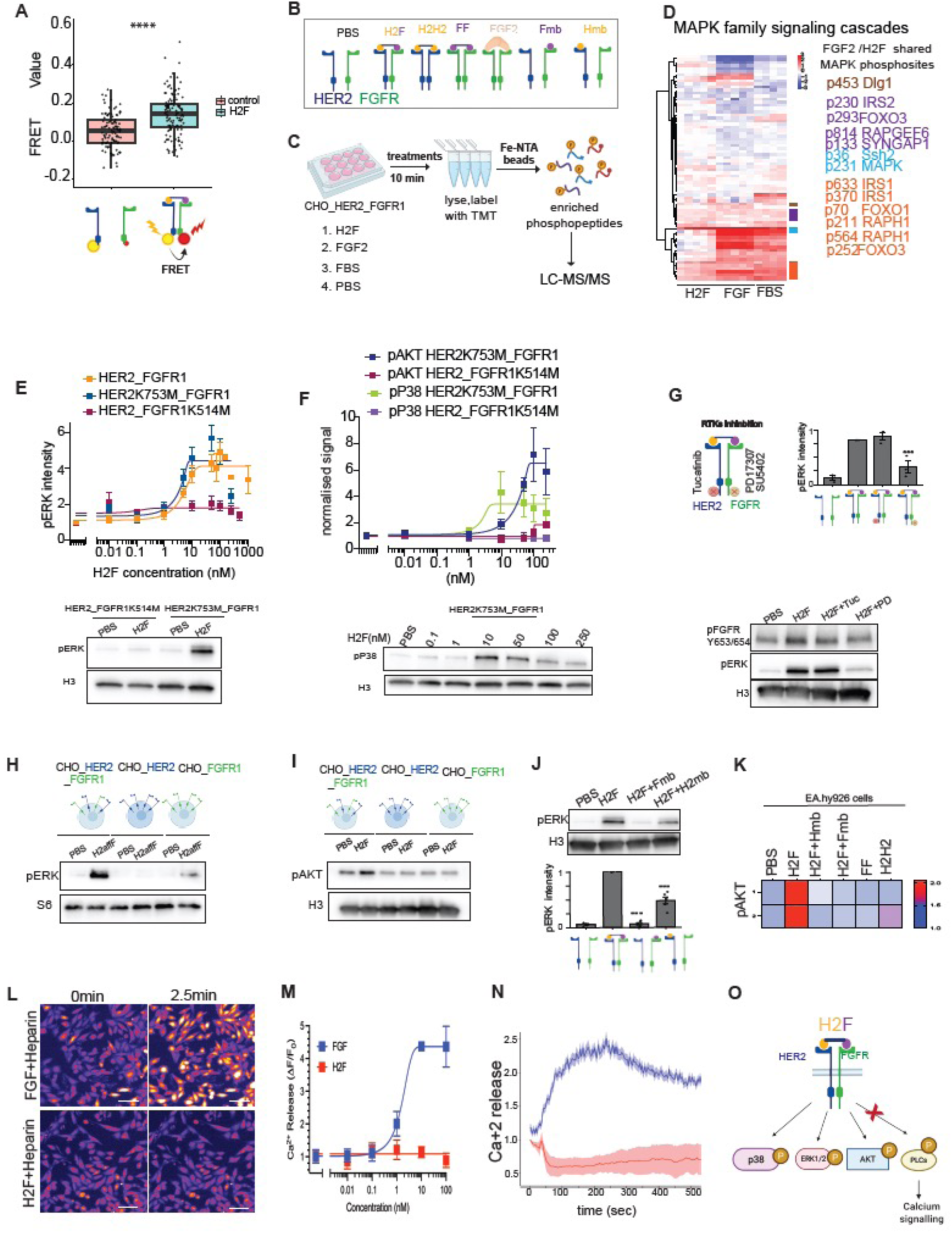
H2F bifurcates RTK mediated MAPK and PLCγ /Ca signaling response. (A) FRET performed in CHO cells expressing human HER2△C-mEYFP and FGFR1△C-mCherry. A scatter plot showing FRET efficiency (Eapp) significantly increased in presence of 500nM H2F in CHO cells expressing human HER2-mEYFP (donor) and FGFR1c-mCherry(acceptor) compared to PBS control. (B) Schematic showing Her2 and FGFR1 homo/heterodimerization in presence of the various H2F, FF, H2H2, FGF2 or competition with Fmb/H2mb.(C) Schematic showing phosphoproteomics flow. (D) MAPK signaling cascade analysis analysed from phosphoproteomics data in 10 min of H2F, FGF2 and FBS treatment in CHO cells expressing human FGFR1c and Human Her2 receptors. Shows list of MAPK phosphosites shared between H2F and FGF2.(E)Graph of H2F titration curve in CHO cells expressing both human HER2_FGFR1c, HER2_FGFR1K514M, HER2K753M_FGFR1. immunoblot showing pERK signal upon 50nM H2F treatment for 10 min in HER2_FGFR1K514M and HER2K753M_FGFR1 expressing CHO cells. (F) Graph of H2F titration curve in CHO cells expressing HER2_FGFR1K514M, HER2K753M_FGFR1 of pAKT and Pp38. Immunoblot showing Pp38 signal upon 50nM H2F treatment for 10 min in HER2_FGFR1K514M and HER2K753M_FGFR1 expressing CHO cells. (G) Model showing selective kinase domain small molecule inhibitors, Tucatinib and PD173074 against HER2 and FGFR respectively. Western blot depicting the effect of PD173074 and Tucatinib treatment on H2F treatment on CHO-hHER2-hFGFR cells FGFR receptor phosphorylation. (H) pERK signal in 100nM H2affF treatment in CHO cells expressing both hHER2 and hFGFR1 and either hHER2 or hFGFR. (I) H2F shows pAKT signals in CHO-hHER2-hFGFR1c cell lines but not in either hHER2 or hFGFR. (J) H2F shows pERK in CHO-hHER2-hFGFR1c cell lines. H2F in competition with either the HER2mb or FGFR1cmb (monomeric minibinders used in excess to block one half of H2F) shows reduced pERK. (K) Heatmap showing fold change in pAKT signal in FACS sorted cells compared to PBS in EA.hy926 cells; competition with either of the Hmb or Fmb reduced the pAKT signal. Homodimers such as H2mb:H2mb or Fmb:Fmb do not show strong pAKT signals. (L) Representative images of timelapse showing Ca^2+^ signaling using Calbryte dye in FGF+Heparin and H2F+heparin treatments. (M) Titration curve of Ca^2+^ signaling using Calbryte dye in FGF+Heparin and H2F+heparin treatments. (N) Quantification of the Ca^2+^ signal intensity shows that FGF+heparin treatment is able to trigger calcium release in CHO-hHER2-hFGFR1c cells, whereas treatment with H2F+heparin is not. (O) Model showing that HER2mb:FGFR1cmb phosphorylates the Y653/654 residues of FGFR, activating the MAPK/pAKT pathways, but fails to show Ca signaling response. Thus, H2F bifurcates RTKs mediated MAPK and PLCγ /Ca signaling response, scale bar= 50um. ***p < 0.001, ****p < 0.0001.

To characterize the phosphoproteomic signatures induced by H2F, we treated CHO cells overexpressing human HER2 and FGFR1 with FGF, H2F, FBS or vehicle (no minibinder), and performed quantitative phosphopeptide enrichment analysis after LC-MS/MS. Across all conditions, 8,077 unique phosphosites were identified, of which 6,325 sites were sufficiently quantified and localized for further analysis (Fig-4SC). Comparative analysis revealed that while H2F and FGF shared a large subset of phosphorylated targets, including canonical MAPK pathway components such as MEK1 and ERK1/2, H2F also induced a unique phosphosignature (Fig. 4C,D). This suggests that H2F induces a partially overlapping but mechanistically distinct signaling program compared to native FGF ligands potentially reflecting altered receptor dimer topology and biased signal transduction downstream of HER2–FGFR1 heterodimers (Fig. 4D,4SD-G).

The dose-response curve for H2F-induced pERK revealed an EC₅₀ of 5.84 nM (Fig. 4E), indicating high potency and specificity. To determine whether the kinase activities of HER2 and FGFR1 were required for H2F-dependent ERK activation, we examined CHO cells coexpressing kinase-null HER2 (HER2_K753M) with wild-type FGFR1, and kinase-null FGFR1 (FGFR1_K514M) with wild-type HER2. H2F failed to induce pERK, pAKT,pP38 in the presence of kinase-null FGFR1, whereas a robust pERK response persisted when HER2 was kinase-null (Fig. 4E,F). In CHO cells coexpressing wild-type HER2 and FGFR1, pretreatment with the HER2-specific kinase inhibitor Tucatinib¹⁵ had no effect on H2F-induced pERK, whereas FGFR inhibitors PD173074 markedly reduced ERK phosphorylation (Fig. 4G). These findings demonstrate that H2F-mediated pERK activation is strictly dependent on FGFR1 kinase activity but independent of HER2 kinase activity.

To further dissect whether proximity of both HER2 and FGFR receptors is needed to trigger downstream MAPK signaling, we examined three novokine constructs (H2F, FH2, and H2affF) designed to bind both HER2 and FGFR1c simultaneously. H2F Novokines act as agonists, inducing robust phosphorylation of ERK, AKT, and p38 MAP kinases exclusively in cells co-expressing both HER2 and FGFR1, whereas cells expressing only HER2 or only FGFR1 showed no activation of these pathways (Fig. 4H,I; Fig. S4I,J). Furthermore, the signaling response was abolished by antagonistic HER2 or FGFR mini-binders (Fig. 4J, 4SI). Similar results were obtained with two independently designed HER2-binding constructs, strongly suggesting that H2F signaling is mediated by proximity-induced heterodimerization of HER2 and FGFR1c. In EA.hy926 cells (an immortalized human umbilical vein endothelial cell line), expressing native FGFR and HER2, treatment with the H2F, FH2mb, or H2affF constructs each induced phosphorylation of AKT (pAKT), whereas competition with either H2mb or Fmb markedly reduced pAKT levels (Fig. 4K, S4O,P). The signaling profile induced by the heterodimers H2F and FH2mb was distinct from that induced by the homodimers H2mb:H2mb (H2H2) and Fmb:Fmb (FF). Neither homodimer (H2H2 or FF) enhanced transdifferentiation efficiency or elicited pAKT signaling (Fig. 4J), in contrast to H2F. Taken together, these findings indicate that both receptors are required for the observed downstream signaling.

### HER2-FGFR novokine represses PLCγ activation and acts via MAPK arm of FGFR

FGF and EGF activate a common set of intracellular signaling proteins (e.g. Shc, FRS2, PLCγ, Grb2, Src, PI3K) upon ligand-induced tyrosine phosphorylation ^11,12^(Fig. S4H). To investigate how H2F-induced HER2– FGFR1 heterodimers alter FGFR signaling, we examined FGFR1 phosphorylation after H2F treatment. H2F induced robust phosphorylation of FGFR1 at the activation loop tyrosines Y653/Y654 – modifications that are essential for FGFR kinase activation– but did not induce phosphorylation of Y766 (Fig.S4L,M,N). FGFR phosphorylation on Y653/654 is also abrogated by FGFR RTKs inhibitors (Fig.4E,S4K). the Phosphorylation of Y766 is required for PLCγ recruitment and downstream Ca²⁺ mobilization^13,14^. Consistent with this requirement, H2F stimulation failed to evoke any intracellular Ca²⁺ transient (monitored by a fluorescent Ca²⁺ indicator^1^), whereas FGF or a C6-79C-mb7(FGFR1/2c designed agonist) agonist triggered a clear Ca²⁺ response (Fig. 4L-N). Meanwhile, the maintained phosphorylation at Y653/Y654 indicates preserved activation of the MAPK cascade.

Thus, H2F-induced HER2–FGFR1 heterodimers selectively propagate MAPK signaling while bypassing the PLCγ/Ca2+ branch, effectively segregating the canonical FGFR outputs (Fig. 4O). Collectively, these findings demonstrate a biased FGFR signaling response upon HER2–FGFR heterodimerization: FGFR kinase– dependent ERK/AKT pathways are engaged while the PLCγ/Ca^2+^ axis is excluded.

### H2F induced biased FGFR signaling supports pluripotency but not endothelial differentiation

The activation of FGFR1/2c by H2F, which notably lacks the Ca²⁺ signaling branch, enables us to pinpoint biological processes that rely on this specific FGFR modality. During vasculogenesis, both endothelial and perivascular cells arise from common mesodermal precursors, and FGF signaling plays a crucial role in their fate determination ^18,19^. Our previous study demonstrated that selective activation of the FGFR1/2c isoform using a designed FGF agonist (C6-79C-mb7) strongly promoted endothelial differentiation over perivascular fate, underscoring FGFR1/2c’s role in endothelial lineage specification^1^. We therefore investigated whether the H2F Novokine, which pairs FGFR1/2c-splice variants with HER2 and activates only a partial FGFR pathway, could influence cell fate in a vascular differentiation model. To test this, we replaced FGF2 in the endothelial differentiation protocol with 100 nM of H2F, H2affF, or the monomeric FGFR1/2c inhibitor mb7 (Fig. 5A). Furthermore, we showed that both FGFR and Her2 were expressed at this stage of differentiation (Fig.S5A). Flow cytometry analysis further showed that 75.48% of FGF2-treated cells expressed the endothelial marker VE-CAD, whereas with H2affF treatment only 21.14% expressed VE-CAD and with H2F treatment only 0.03% expressed VE-CAD (Fig. 5B). Immunofluorescence imaging revealed that treatment with H2F and H2affF predominantly induced expression of the perivascular marker PDGFR-β (Fig. 5C). These findings suggest that Novokines, by pairing FGFR with HER2, disrupt FGF-mediated endothelial differentiation and instead bias cell fate toward the perivascular lineage.

**Figure 5:**
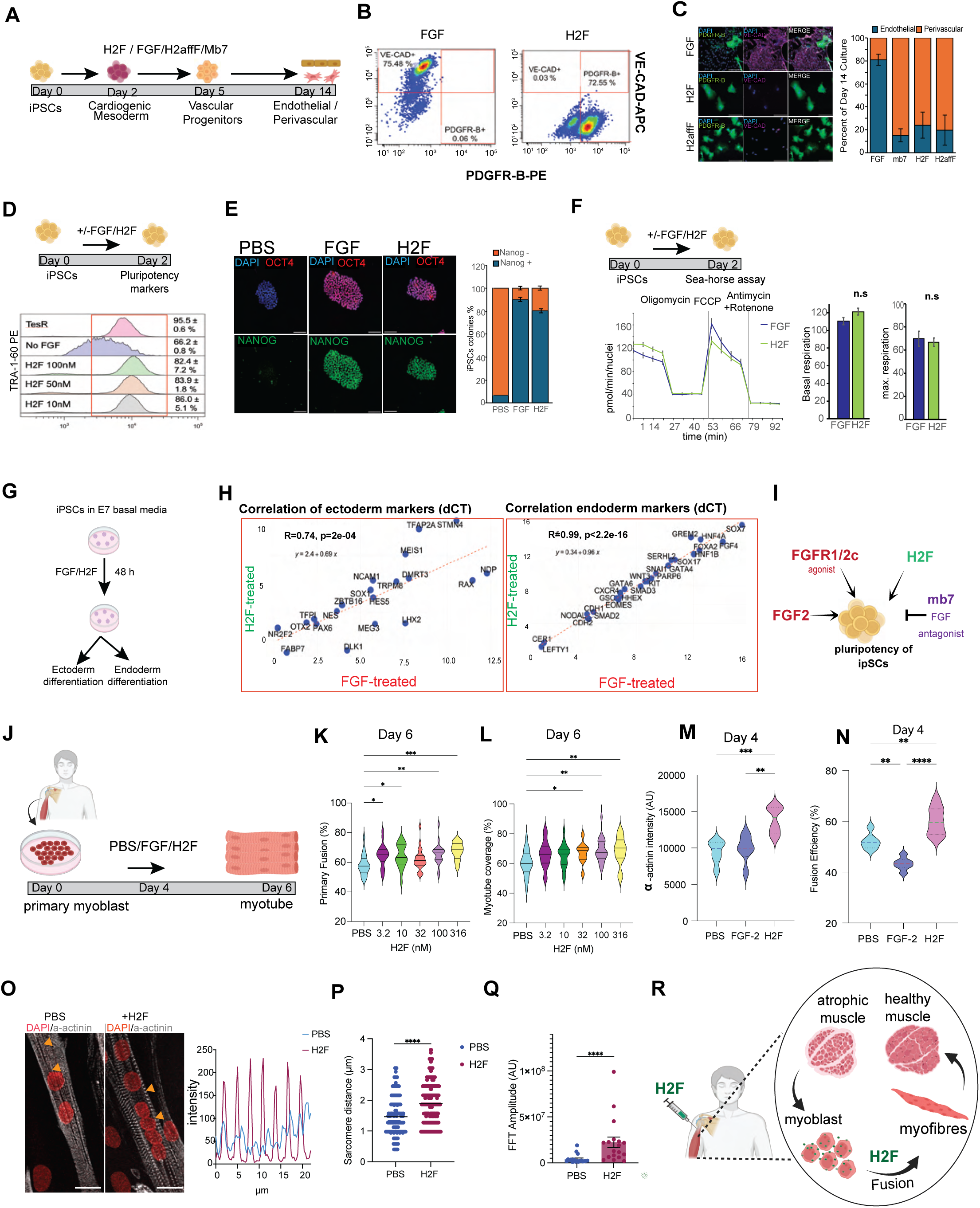
Effects of H2F on endothelial differentiation, pluripotency maintenance in iPSCs and primary myoblast differentiation. (A) Schematic of the differentiation protocol from iPSCs to endothelial and perivascular lineages, highlighting intermediates. (B) Proportion of endothelial or perivascular cells generated at day 14 following treatment with FGF2, mb7, H2F. Error bars represent 2 independent biological repeats. Representative scatterplots of endothelial (VE-cadherin) and perivascular (PDGFR-B) markers, assayed using flow cytometry. (C)Immunofluorescence images showing PDGFR-B and VE-CAD expression at day 14 in cultures treated with FGF, H2F, or H2affiF. Scale bar: 100um. (D) Schematic of experimental setup to assess pluripotency marker expression in iPSCs after 48-hour treatment with FGF or H2F. Representative histograms of TRA-1-60 expression in iPSCs following treatment with varying concentrations of H2F, assayed using flow cytometry. Error represents 2 independent biological repeats. (E) Immunofluorescence images showing Oct4 and Nanog expression in iPSCs treated with FGF or H2F for 48 hours. Scale bar: 100um. (F) Oxygen consumption rate (OCR) across time for iPSCs treated with FGF or H2F. Timepoints indicate the addition of mitochondrial inhibitors (oligomycin; FCCP; antimycin A/rotenone). Oxygen consumption rate graph of 48 hours H2F and FGF treated iPSCs in E8 base media. Bar graphs show maximal respiration, spare respiratory capacity, and ATP Production. Graphs show no significant difference between H2F and FGF. (G) A schematic showing iPSCs treated with FGF/H2F in minimal media for 48h followed by Ectoderm/endoderm differentiation and RNA isolation. (H) H2F treated and FGF treated iPSCs undergo ectoderm and endoderm differentiation efficiently shown by a positive correlation of ectoderm markers (left graph) and endoderm marker expression (right graph) in respective differentiations. (I) Model summarizing the effects of H2F on stem cell fate: Biased FGFRc activation maintains pluripotency, while its inhibition drives differentiation. (J) Schematic showing patient derived myoblast differentiation to Myotubes. (K)Fusion efficiency of primary myoblasts in untreated and various concentrations of H2F. (L) Area coverage of primary myoblasts in untreated and various concentrations of H2F. (M)Alpha-actinin intensity quantification in PBS,10nM FGF2 treated versus 100nM H2F is compared at day 4.(N)Fusion efficiency of primary myoblasts in PBS, 10nM FGF versus 100nM H2F at day 4. (O)Immunofluorescence images showing untreated and 100nM H2F treated primary myotubes at day 6, stained for a-actinin(gray) and DAPI (blue), scale bar= 20um. A small yellow triangle marks the length measured in the plot profile. The graph shows the sarcomere distribution along the myofibrils in PBS versus H2F.(P) Distance between z-discs is measured in PBS versus 100nM H2F treated myofibers at day 6. N=4, PBS n=230, H2F n=453. (Q) Quantification of Z-disk signal clarity, measured as the FFT Peak Amplitude. (R) A model showing patient derived primary myoblast can be differentiated efficiently into muscle cells with potential of muscle atrophies treatment. ****p < 0.0001, * p < 0.05

Since FGFR signaling is essential for human stem cell pluripotency ^20,21^, we aimed to determine whether H2F-mediated biased FGFR1/2c signaling, which selectively activates the MAPK pathway while bypassing the Ca²⁺/PLCγ axis, is sufficient to sustain pluripotency. First, to test if the FGFR1/2c splice variant was sufficient for pluripotency, we treated hiPSCs with the FGFR1/2c-specific antagonist mb7 (100 nM). This led to loss of Oct4 and downregulation of pluripotency markers (Fig. S5B), despite active b-isoforms, indicating FGFR1/2c is essential for pluripotency. Conversely, replacing FGF with the FGFR1/2c-specific agonist C6-79C-mb7 maintained 97–99% TRA-1-60⁺ cells (Fig. S5C), outperforming FGF. These results demonstrate that FGFR1/2c splice variant activation is both necessary and sufficient for maintaining human stem cell pluripotency. To further test H2F biased FGFR1c signaling in pluripotency, we treated WTC11 hiPSCs with C6-79C-mb7, mb7, FGF, or H2F in minimal media (E8) for 48 hours (Fig. 5D). Flow cytometry and immunostaining analyses showed that cells treated with C6-79C-mb7, FGF, or H2F maintained the expression of pluripotency transcription factors (OCT4, Nanog) and the surface markers TRA1-60/TRA1-81, whereas mb7-treated cells and FGF-deprived controls showed reduced marker expression (Fig. 5D, E, S5C). Seahorse analysis indicated that H2F-induced FGFR signaling did not alter mitochondrial respiration compared to the activation seen with FGF2 (Fig. 5F). The H2F treated iPSCs showed a positive correlation with FGF treated iPSCs for ectoderm and endoderm lineage differentiation (Fig-5G,H). These findings suggest that H2F-mediated signaling is sufficient to sustain hiPSC pluripotency, while FGFR dependent Ca²⁺/PLCγ signaling is dispensable (Fig. 5I). FGF2 is a known component of stem cell maintenance media (e.g., mTeSR); our results indicate that specifically the FGFR1c/2c–MAPK axis can uphold the pluripotency.

### H2F enhances primary Myoblast fusion and maturation

In primary myoblasts from adult patients, H2F treatment significantly increased fusion efficiency and myotube coverage area compared to FGF treated controls, yielding thicker, more continuous fibers (Fig. 5J-N, S5F). Comparable effects were also observed in human iPSC-derived myoblasts (Fig. 5SD). Unlike native FGF, which primarily drives myoblast proliferation, H2F elevated both fusion efficiency and α-actinin expression (Fig. 5M-N, S5E), underscoring a distinct mechanism of action. Sarcomere analysis further revealed that H2F promotes myotube maturation, with well-defined striations and physiological range for Z-disc spacing (median 1.87 µm) compared to the disorganized sarcomeres of the PBS-treated controls (median 1.28 µm) (Fig.5O-Q). Importantly, these results demonstrate that H2F is not limited to fetal or iPSC-derived myogenic cells; it is equally effective in adult primary myoblasts, where regenerative potential is typically diminished. Thus, H2F emerges as a promising priming agent for autologous myoblast transplantation and a potential therapeutic to enhance muscle regeneration in settings of atrophy, injury, or aging (Fig. 5R).

## Discussion

We find that pairing HER2 and the FGFR1/2c using the Novokine H2F drives fibroblast-to-muscle reprogramming. Similar results were obtained with H2F Novokine-constructs carrying two distinct HER2 binding domains, strongly supporting that the observed activity reflects HER2 recruitment. FRET analysis confirmed that H2F mediates HER2-FGFR heterodimerization, a critical step in reprogramming. Phospho-proteomic profiling revealed that H2F engages signaling nodes such as various GAPs, GEFs, and MAPKs, overlapping partially with canonical FGF signaling. Mechanistic dissection showed that HER2-FGFR heterodimerization selectively bifurcates RTK signaling, activating MAPK/AKT while bypassing PLCγ. H2F relies on canonical FGFR Y653/Y654 phosphorylation for downstream signaling. By activating the MAPK branch while avoiding Ca²⁺-dependent signaling, H2F enables dissection of FGFR1/2c-dependent pathways across diverse biological context.

H2F-mediated FGFR activation is incompatible with endothelial differentiation, but crucial in myogenic differentiation, myofiber maturation, and pluripotency maintenance in human stem cells, suggesting that FGFR-dependent Ca²⁺ signaling is dispensable for these processes. Single-cell RNA-seq from the Tabula Sapiens atlas identified human cell types co-expressing HER2 and FGFR1/2c, pointing to potential in vivo contexts for H2F activity (Fig. S5G,H). As proof of concept, patient-derived primary myoblasts differentiated efficiently with H2F treatment, underscoring its promise for muscle regenerative medicine.

Our strategy of fusing designed protein domains to recruit target receptor pairs should be broadly applicable. As described in Abedi et al., we have used similar approaches to generate novel cytokine-like molecules that pair new combinations of cytokine receptors. While previous studies with orthogonal receptor systems have demonstrated that unrelated cytoplasmic domains can initiate signaling 32, our designed Novokines now pioneer and establish the utility of the approach in natural cellular context. Novokines harness the ability to bring endogenously existing receptors together, creating unprecedented diversity of the receptor combinations and targeted cell types. With the rapidly expanding set of AI-designed binding domains, libraries of Novokines can be systematically generated and screened across reprogramming and differentiation assays. As demonstrated with H2F, such synthetic ligands hold dual promise: they can serve as therapeutic leads for regenerative medicine and simultaneously uncover foundational principles of receptor wiring and signaling network logic.

## Resources

scRNAseq: https://doi.org/10.7910/DVN/2B5OGU bulk seq: https://doi.org/10.7910/DVN/C42ZWI

## Acknowledgements

We thank Dale Hailey and the Garvey Microscopy Core for assistance with microscopy, and Mary C. Regier and the UW Genomics Core for guidance with RNA sequencing. We are grateful to Prof. Adam Gebbale (Fred Hutchinson Cancer Center) for providing the HER2-specific kinase inhibitor Tucatinib, and to Prof. J. Schlessinger (Yale School of Medicine) for insightful comments on this study. We thank Dr. Shiri Levy, Ethan Narog, Melodie Chiu, Qigong Li and other members of the HRB lab. We thank Mateusz, Johns Hopkins for insights on receptor proximity.

## Materials and Methods

### Cell culture-CHO cells, immortalized HUVEC cells, HFF-MyoD, Primary human myoblasts IPSCs

HFF-MyoD cells-HFF-MyoD cells were maintained in growth media (DMEM with 4.5g/L glucose and Glutamax [Gibco #10566024], 10% fetal bovine serum (FBS) [BioWest #S1620], 1% penicillin/streptomycin (P/S) [Gibco #15140122], 1% NEAA [Gibco #11140050]) at 37°C and 5% CO2. At 80% confluency HFF-MyoD cells were passaged and seeded at a 1:10 ratio. Chinese hamster ovary (CHO) cells - CHO cells were maintained in CHO growth media (F12K [ATCC #30-2004; Gibco #21127022], 10% FBS, 1% P/S) at 37°C and 5% CO2. At 90% confluency CHO cells were passaged and seeded at a 1:10 ratio. Continued background selection of CHO overexpression (OE) lines was maintained through addition of 5 μg/mL blasticidin [Company, catalog no] or 10 μg/ml puromycin [Gibco #A1113803] to the CHO growth media depending on the plasmid used. Immortalized human umbilical vein endothelial cells (HUVECs) - EA.hy926 cells were maintained in growth media (DMEM with 4.5g/L glucose and Glutamax, 10% FBS, 1% P/S, 1% NEAA) at 37°C and 5% CO2. At 80% confluency EA.hy926 cells were passaged and seeded at a 1:10 ratio.Primary human skeletal muscle myoblasts [Cook Myosite #SK-1111] and IPSC derived myoblasts [27] were maintained in Skeletal muscle growth media (SKGM) consisting of 10% FBS [Hycone #SH30071.03), 100nM Insulin [Lonza #BE02-033E20], 40ng/mL FGF-2[R&D Systems #3718-FB] & 1µM Dexamethonsone [Sigma #D4902] in DMEM with 4.5g/L glucose and Glutamax [Gibco #10566024].

### Generation of Skeletal Muscle Myoblasts from iPSCs

iPSC-derived myoblasts (iPSC-MBs) were generated from the UC3-4 iPSC line using a multi-stage protocol adapted from previous work [27]. iPSCs were plated at 15,000 cells/cm² in mTeSR Plus (Stem Cell Tech) and grown to ∼40% confluency in 6-well plates. Myogenic induction was initiated (day 0) with differentiation medium containing DMEM/F12 (Gibco #10565018), 1x non-essential amino acids (Gibco #11140050), 1x Insulin-Transferrin-Selenium (Gibco #41400045), 3 µM CHIR99021 (Axon #1386), and 0.2 µg/mL LDN193189 (Miltenyi Biotec #130-106-540). On day 2, medium was supplemented with 20 ng/mL bFGF (R&D Systems #3718-FB). On day 5, CHIR99021 and ITS were removed and 15% KSR (Gibco #10828028), 2 ng/mL IGF-1 (R&D Systems #291-G1), and 10 ng/mL HGF (R&D Systems #294-HGN) were added. On day 7, bFGF and LDN193189 were removed. Medium was changed every other day until day 28. At day 28 post-induction, a single confluent cell sheet was mechanically dissociated into small clumps and replated onto a Matrigel-coated T225 flask for expansion in SKGM, with feeding every 2–3 days. At day 32, cells were lifted, filtered through a 40µm filter and subjected to fluorescence-activated cell sorting (FACS) to purify for positive Nerve derived growth factor (NGFR+) population [Biolegend #345108]. All downstream experiments used cells between passages 4 and 5.

### Screening novokines in Myogenic transdifferentiation assay

In a 96 well plate, 10^3 per well HFF-tet-on inducible MyoD cells in DMEM with 10% FBS were seeded. Post 2 days of seeding, for the next 4 days cells were treated with 2 nM doxycycline [company, catalog number] in SMM media. The following 4 days cells are incubated with 100nM of each novokine. Cells were fixed in 4% paraformaldehyde [Electron Microscopy Sciences #15710, diluted in phosphate buffered saline (PBS)] for 15 min, washed with PBS, and stained for Desmin/MHC, followed by imaging at Leica spinning disk microscope, stitched (4x4) images of each 96 well at 10X was captured. In Fiji ImageJ, the desmin intensity area was thresholded and the intensity was measured as the desmin positive area.

### Screening myogenic fusion of Primary and IPSC derived myoblasts

Primary and iPSC-derived myoblasts were seeded In 24 well plate polymer bottom plates [Cellvis #P24-1.5P] at 15,000/cm² and grown in SKGM until confluent. To initiate myogenic differentiation (day 0), media was changed to DMEM high glucose supplement with 2% Horse serum [Gibco #16050114] +/- H2F of varying concentrations. The media was changed every 48 hours. On Day 6, cultures were fixed in 4% paraformaldehyde for 20 minutes at room temperature. Cells were then permeabilized and blocked for 1 hour using a blocking buffer consisting of 0.2% Triton X-100, 1% BSA, and 5% Donkey Serum in PBS. Following blocking, cells were incubated overnight at 4°C with a primary antibody against sarcomeric α-actinin (R&D Systems, #MAB9830, at 1:500 dilution. After washing, cells were incubated with a fluorescently-labeled secondary antibody and counterstained with DAPI to visualize nuclei. Images were acquired on a Yokogawa W1 spinning disk confocal microscope at 20x magnification. For each well, a large 4x4 image panel was captured to provide a comprehensive view of the culture. An in-house ImageJ macro was used for automated, unbiased quantification. First, the LABKIT plugin [28] was used to segment a myotube mask from the ɑ-actinin channel. Next, the StarDist plugin [29] was used to label the nuclei from the DAPI channel. A nucleus was classified as “fused” if it met two criteria: (1) area > 40 µm², and (2) >90% of its area overlapped with the myotube mask. Fusion efficiency was calculated as the percentage of fused nuclei relative to the total nuclei per image.

### Design of HER2 minibinder

The top 96 hits from sequencing were expressed and purified from 1 mL E. coli cultures in 96 well plate format, followed by screening on the Octet/BLI system using a two-point titration (1000 nM and 200 nM). The top 10 binders were selected based on binder saturation at both concentrations as well as the flatness of dissociation traces observed. For these 10 binders, we performed a three-fold titration series from 1000 nM to 4 nM, which demonstrated low picomolar affinities via global kinetic fitting (Supplementary Figure1**B**). However, due to the small size (6 kDa) of the binders, the signals were relatively small when binding to the much larger, glycosylated HER2 target protein (70-90 kDa). To address this, the top 10 binders were expressed as GFP fusion proteins (∼33 kDa), purified from 50 mL E. coli cultures, and reassessed using Octet/BLI to enhance the signal-to-noise ratio. The GFP fusion proteins produced significantly larger signals, consistent with the original kinetic fitting results, confirming low pM affinities with near-flat dissociation rates over the duration of the experiment (Figure1C). The top binder exhibited a single, monodisperse peak in size exclusion chromatography (SEC), both with and without the GFP fusion tag (S1A). Structural analysis of the design model revealed key interactions with HER2, where PHE51 and TYR32 from the minibinder fit into complementary hydrophobic pockets of the target (Figure1C).

### Ca^2+^ assay

CHO-hHER2-hFGFR1c cells were seeded on 96-well flat bottom microplates [Corning #3603] and grew to 70-80% confluence. Cells were starved in serum-free F12-K medium for 3 hours. Following starvation, the cells were incubated in serum-free media containing 5mM Calbryte 520 AM fluorescent intracellular calcium indicator [AAT Bioquest, #20651] for 30 min at 37°C. Cells were washed 3X with serum-free media and treated with various concentrations of recombinant FGF2 (with or without 40mg/mL Heparin [Iduron, #H010]) or designed scaffolds. Confocal live imaging was done on a Nikon Yokogawa W1 spinning disk confocal microscope using a 20X objective. Parameters for each live frame: Excitation/Emission filters for GFP fluorescence, Exposure time of 150ms, Acquisition rate of 5 sec/frame, and total recording time of 20 minutes (5 min baseline recording + 15 min ligand treatment time). Images were processed with Fiji software distribution of ImageJ v1.52i84,93 and frame-by-frame cellular fluorescence intensity was tracked and quantified with CellProfiler.94,95 Dose-specific average calcium release was calculated by tracking each individual cell’s response during the recording time and computing the mean peak fluorescence achieved by all cells in the frame. An average of 50-100 cells were tracked per recording.

### In vitro differentiation of endothelial cells

Briefly, hiPSCs (WTC-11 human induced pluripotent stem cells) [Coriell, #GM25256] were seeded on 24-well plates coated with growth factor-reduced Matrigel [Corning, #356231] and cultured in mTeSR1 stem cell medium [StemCell Technologies, #85850] until cells reach confluence with media changes daily. One day before differentiation (deemed Day (-1)), cells were pre-treated with mTeSR1 supplemented with 1µM of GSK3-Inhibitor (CHIR99021) [Cayman Chemicals, #13122]. On the first day of differentiation (D0), stem cell media was replaced with cardiogenic mesoderm media consisting of RPMI 1640 Medium [Thermo, #11875093] supplemented with 1X B27(-) [Fisher Scientific, #A1895601], 100 ng/mL Activin A [PeproTech, #120-14P] and 1X Matrigel for 17 hrs. The next day, media was replaced with RPMI supplemented with 1µM of GSK3-Inhibitor (CHIR99021), B27 (-), and 5 ng/mL bone morphogenetic protein-4 (BMP-4) [R&D systems, #314-BP-010] for 24 hours. On Day 2 of differentiation, cells were washed with 1X PBS and media was replaced with vascular differentiation media consisting of StemPro [Thermo Fisher, #10639011] supplemented with 1X Glutamax, 1X Penicillin-Streptomycin, 300 ng/mL vascular endothelial growth factor (VEGF) [R&D systems, #293-VE-050], 10 ng/mL BMP-4, 5 ng/mL FGF2, 50 ug/mL Ascorbic Acid [Sigma-Aldrich, #A8960], and 40 µM monothioglycerol (MTG) [Sigma-Aldrich, #M6145]. On Day 5, cells were dissociated with Accutase [Thermo, #A1110501] and replated on 12-well 0.1% gelatin-coated tissue culture dishes in endothelial growth media (EGM) consisting of EGM basal media [Lonza, #CC-3121] supplemented with 20 ng/mL VEGF, 20 ng/mL FGF2 and 1µM GSK3-Inhibitor (CHIR99021). EGM media was replaced every 48 hours until the final harvest at Day 14.

### Flow cytometry

Cells derived using FGF2, H2F, and mb7 (at Day 14 of differentiation) were harvested as a single cell suspension, adjusted to a concentration of 0.5 x 10^6^ cells/mL in ice-cold FACS Buffer (1X PBS, 1% BSA, 0.1% sodium azide [Millipore-Sigma, #26628-22-8]), and blocked for 30 minutes on ice. Cells were washed 3 times in 1X PBS by centrifugation at 1500 rpm for 5 minutes each, following which primary (labeled) antibodies were added at a dilution of 1:100 in FACS buffer for 1 hr at 4C in the dark: VE-Cadherin-APC [eBioscience, #17-1449-42], PDGFR-B-APC [BioLegend, #323608]. Cells were washed 3 times in 1X PBS by centrifugation at 1500 rpm for 5 minutes each and resuspended in 100uL of ice-cold FACS Buffer for subsequent analysis. Unstained controls were included. Samples were run on a FACSCanto II flow cytometer (BD Biosciences) and recorded events were analyzed using the flowCore package for R. FSC and SSC (Unstained control) were used for size gating. Events were analyzed as the percentage of cells positive for the given panel of markers.

### Immunostaining of differentiated iPSCs

For immunofluorescence imaging of differentiated iPSCs, cells were seeded on glass coverslips coated with 0.1% gelatin on Day 5, and cultured until confluency on Day 14 following the process described above. The cells were then fixed in 4% paraformaldehyde (PFA) for 15 minutes. The fixed cells were washed three times for 5 min each in 1X PBS before blocking for 1 hr with 3% BSA and 0.1% Triton X-100 in 1X PBS while on nutation. Primary antibody incubation was carried out at a 1:100 dilution in blocking buffer overnight at 4C: CD31 (Cell Signaling, Catalog #3528), and PDGFR-B (Cell Signaling, Catalog #3169). Following overnight incubation, the cells were washed three times for 5 min each in 1X PBS while on nutation. The cells were then incubated with secondary antibodies (Invitrogen, A21050 and Invitrogen, A11008; 1:200 each) diluted in blocking buffer for 1.5 hrs at 37°C. Secondary antibodies were then removed, and cells were washed three times for 10 min each in 1X PBS on nutation. Coverslips were sealed using VECTASHIELD with DAPI [Vector laboratories, #H-2000-2] upside-down on glass slides for analysis in confocal microscopy. Images were taken on a Nikon Yokogawa W1 spinning disk confocal microscope using a 20X objective.

### Pluripotency assays

hiPSCs (WTC-11 human induced pluripotent stem cells) [Coriell, #GM25256] were seeded onto 12-well plates pre-coated with growth factor-reduced Matrigel [Corning, #356231] at a density of 20,000 cells per well, in mTeSR1 medium supplemented with 10 µM Rock inhibitor (Y-27632, Tocris Bioscience). The following day, the media was replaced with E8 basal media consisting of DMEM/F12 supplemented with L-ascorbic acid-2-phosphate magnesium (64 mg/L), sodium selenite (14 µg/L), insulin (19.4 mg/L), transferrin (10.7 mg/L), FGF2 (100 µg/L), and TGF-β1 (2 µg/L) [33]. For experimental conditions, FGF2 was replaced with 100 nM of the FGFR1c-specific inhibitor minibinder (mb7) or the FGFR-HER2 novokine (H2F). Cells were cultured in these experimental conditions for 48 hours before harvesting for analysis.

### FRET imaging and data analysis

FRET imaging was then performed using a Leica SP8 confocal microscope as discussed in detail previously. The donor scan excited YFP at 488 nm and its emission was collected between 500 and 540 nm. The acceptor scan excited the acceptor at 552 nm, and its emission was collected from 590 to 700 nm. For the FRET scan, YFP was excited, and mCherry emission was detected in the 590 to 700 nm range. Cells were imaged in the absence and in the presence of 500 nM of H2F protein. One or two small (∼5 mm) regions of the plasma membrane were selected per cell for FRET analysis. The FRET efficiency, the donor concentration (HER-YFP), and acceptor concentration (FGFR1c-mCherry) were measured in each region. For both conditions, with and without H2F protein, the FRET efficiency as a function of the acceptor concentration, the donor concentration as a function of the acceptor concentration, and the deviation from the proximity for each point were plotted using the software package Origin. The deviation from proximity FRET (Fig. 4A) was performed using an unpaired t-test with the Prism software.

### Phosphoproteomics Analysis

CHO-hHER2-hFGFR1c cells were seeded on 6-well plates and grown to 80% confluence and serum starved for four hours prior to treatment. Cells were treated for 15 minutes with either 100nM of the FGFR-HER2 novokine, 1 nM of FGF, media containing 10% fetal bovine serum, or serum-free media with PBS. After treatment, cells were washed twice with PBS and lysis buffer (8M urea, 50mM NaCl, 200 mM EPPS [pH 8.5], Roche phosSTOP phosphatase inhibitor tablets, and Roche protease inhibitor tablets) was added lyse cells prior to processing. For phosphoproteomics, 100 μg of protein lysate for each sample was processed. Lysates were reduced (5 mM TCEP [Sigma Aldrich], 20 min) and alkylated (10 mM iodoacetamide [Sigma Aldrich]); excess iodoacetamide was quenched with the addition of 10 mM dithiothreitol [Sigma Aldrich]. Proteins were extracted using the SP3 protocol [34] prior to resuspension in 200 mM EPPS (pH 8.5) and digestion with LysC Wako) for 18 hours with vortexing followed by trypsin digestion for 6 hours at 37°C at 200 rpm. Peptides were labeled with TMTpro reagents [Thermo Fisher Scientific] for 1 hour at room temperature on the SP3 beads, then quenched with 5% hydroxylamine and incubation for 15 minutes at room temperature. TMTpro-labeled peptides were cleaned using C18 SepPaks (Waters) and the resulting eluate was dried to completion in a speedvac. Phosphopeptide enrichment was performed using Fe-NTA magnetic beads (PureCube Fe-NTA MagBeads). Phosphopeptides were cleaned using C18 STAGE-tips [35] and eluates were dried to completion and frozen at −80°C prior to analysis by LC-MS/MS.

Phosphopeptides were resuspended in 5% acetonitrile and 2% formic acid then analyzed using an SPS-MS3 method on an Orbitrap Tribrid Eclipse and a 180 minute method with a flow rate of 300 nl/min on a C18 analytical column (15 cm Odyssey, ionOpticks). A gradient was applied starting in 100% solvent A (0.125% formic acid), then increasing from 4% solvent B (95% acetonitrile, 0.125% formic acid) to 20% solvent B after 165 minutes. MS1 spectra (Orbitrap Resolution = 120,000; maximum injection time = 50 ms; normalized AGC = 200%; RF lens % = 30) were collected for precursor detection cycling between three FAIMS CVs (−40, −60, −80) . MS2 spectra were collected for precursors between 300-1500 m/z using quadrupole isolation, dynamic exclusion (time delay = 90 seconds, window = 10 ppm) and intensity filtering for ion trap spectra (>5000). Ion trap MS2 spectra (CID energy % = 35; multistage activation = true; neutral loss mass = 97.9763; maximum injection time = 35 ms; normalized AGC % = 250) were used for peptide identification. Orbitrap MS3 spectra (HCD energy % = 45; first m/z = 110; Orbitrap Resolution = 50,000; maximum injection time = 86 ms; normalized AGC % = 250) were used for TMTpro reporter ion quantification. Phosphopeptides were identified using the Comet search algorithm and filtered to a 1% protein and peptide false discovery rate using linear discriminant analysis and the rules of protein parsimony.

### Statistical analysis

Two-sided Student’s t-test was performed to determine p-value. For phosphoproteomics analyses, Welch’s t-tests were used to assess statistical significance followed by Benjamini-Hochberg procedure [36] to correct for multiple hypothesis testing.

---

**Supplemental Figure 1.**
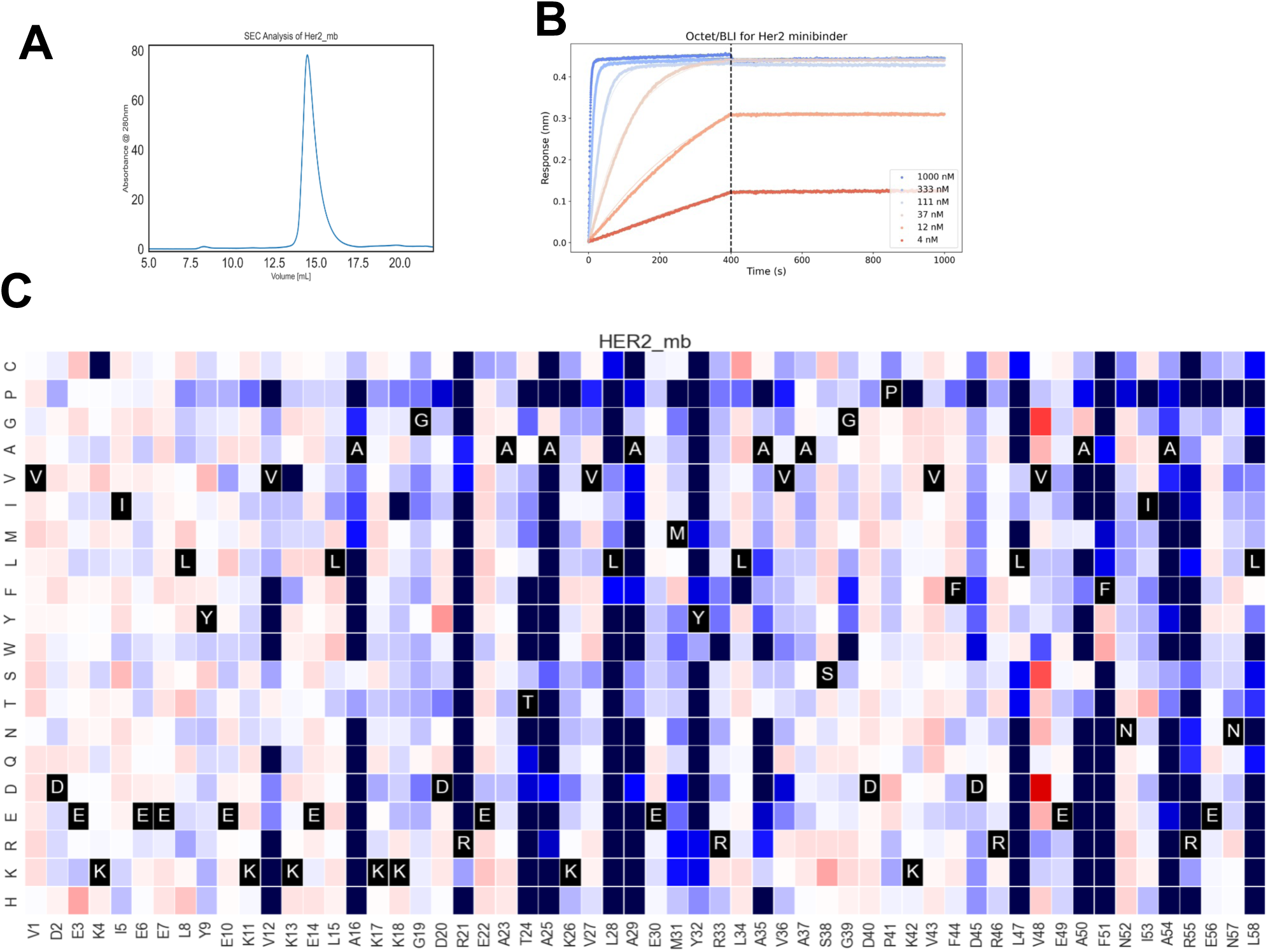
HER2 minibinder. A. Size exclusion chromatography on S75 column of her2 minibinder after IMAC from 50 mL purification shows a monodisperse peak. B. Octet/BLI analysis of Her2 minibinder fused to GFP with three-fold titration from 1000 nM to 4 nM. The binder has low pM Kd based on global kinetic fit, but the off rate is too flat for accurate measurement. C. The SSM for the full binder is shown as well with the color representing yeast surface display binding affinity. The SSM data is additionally shown for surface only residues and core only residues showing the general conservation of core residues and general lack of conservation of surface positions as expected.

**Supplemental Figure 2.**
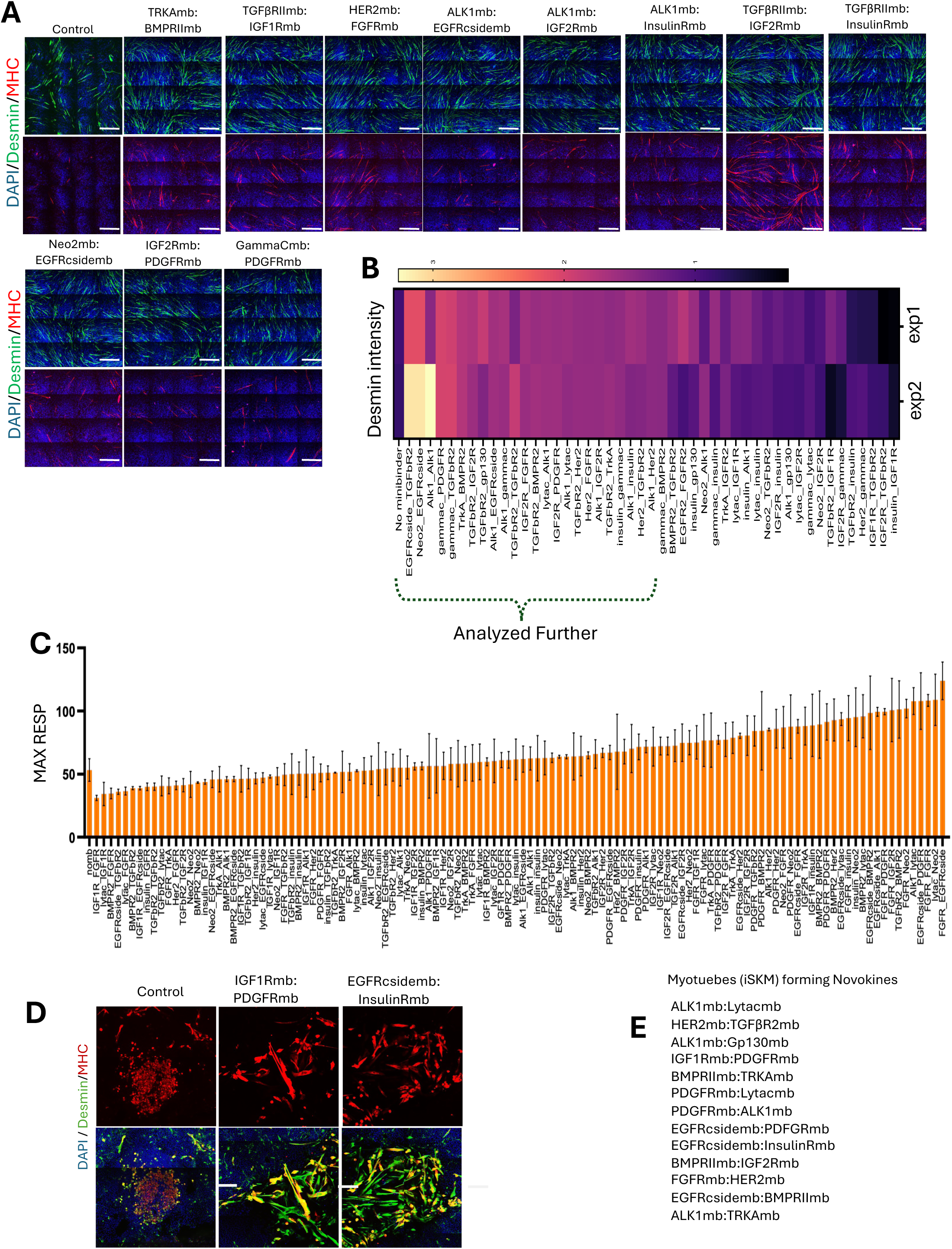
**Designed novokines enhance skeletal muscle reprogramming.** A. Confocal images showing desmin(green), MHC(red) and DAPI(blue) staining of tSKMs from select novokine treatments that increased myogenic trans-differentiation efficiency in assay compared to control. Scale bar=500um B. Heatmap showing myotube desmin intensity compared to the control in two independent secondary screens of designed novokines in the myogenic trans-differentiation assay. Select novokines were screened for activity in flow cytometry phospho-effector screens. C. Graph showing the maximum respiration of control and novokine treated myogenic trans-differentiation assay cells in Seahorse XF Cell Mito Stress Test assay. D. Confocal images showing desmin(green), MHC(red) and DAPI(blue) staining of iSKMs from select novokines treatments that increased iPSCs derived myogenic differentiation efficiency in assay compared to control. Scale bar=100um E. List of the novokines that resulted in the formation of desmin positive iSKM in the myogenic differentiation assay.

**Supplemental Figure 3.**
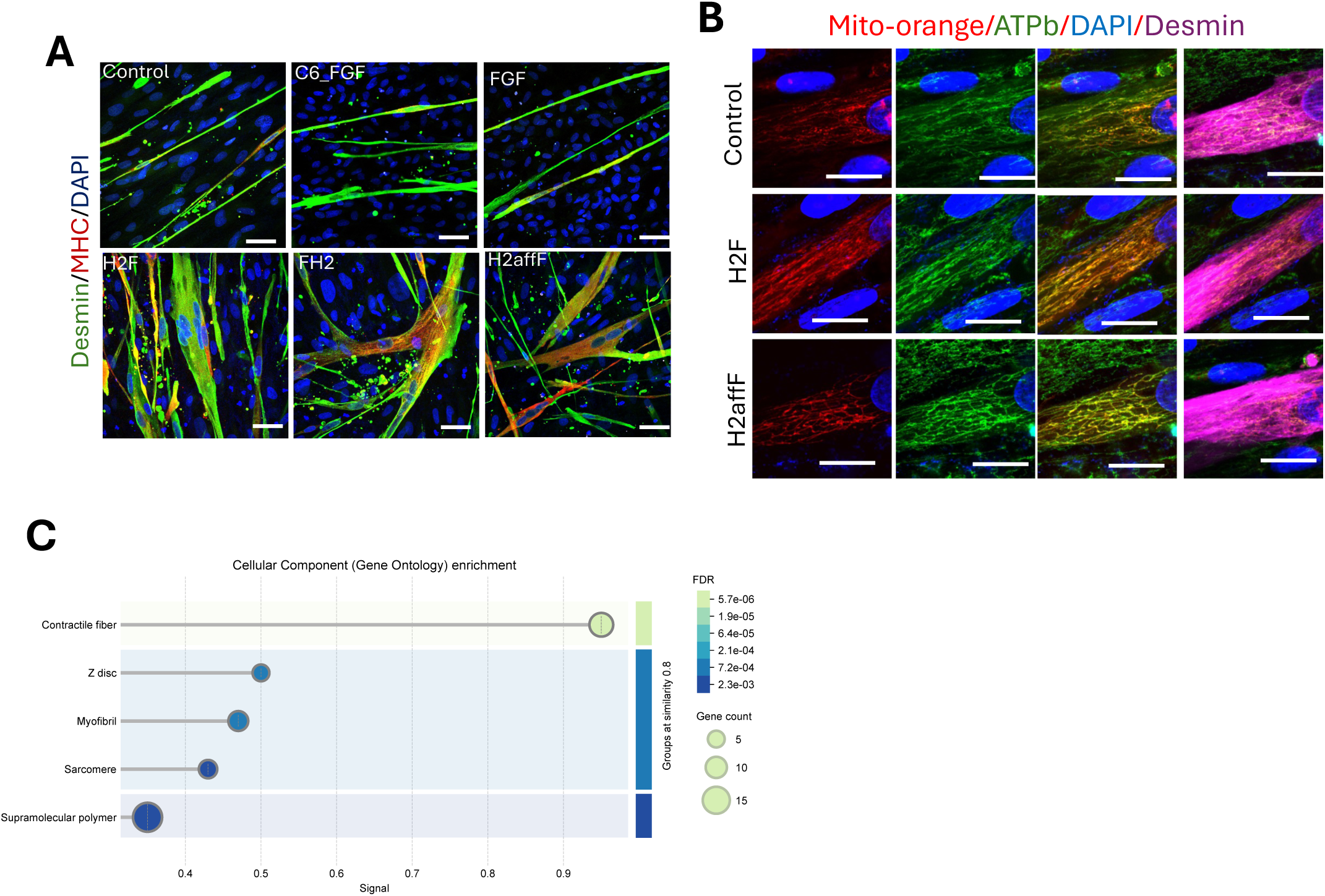
HER2 and FGFR1/2c heterodimerization shows enhanced myogenic reprogramming. A. Confocal image showing desmin staining(green), MHC(red) and DAPI(blue) in control, C6-79C-mb7, FGF, H2F, FH2, and H2affF treatment during myogenic trans-differentiation, scale bar=40um. B. Confocal images of myotubes stained for Mito-Tracker Orange (red), ATPb-Synthase (green), DAPI (blue), and Desmin (magenta) following control, H2F, or H2affF treatment during myogenic trans-differentiation, scale bar=15um.. C. In H2F tSKM versus no mb tSKM bulk mRNA analysis shows that sarcomeric components expression is relatively increased in the H2F treatment.

**Supplemental Figure 4.**
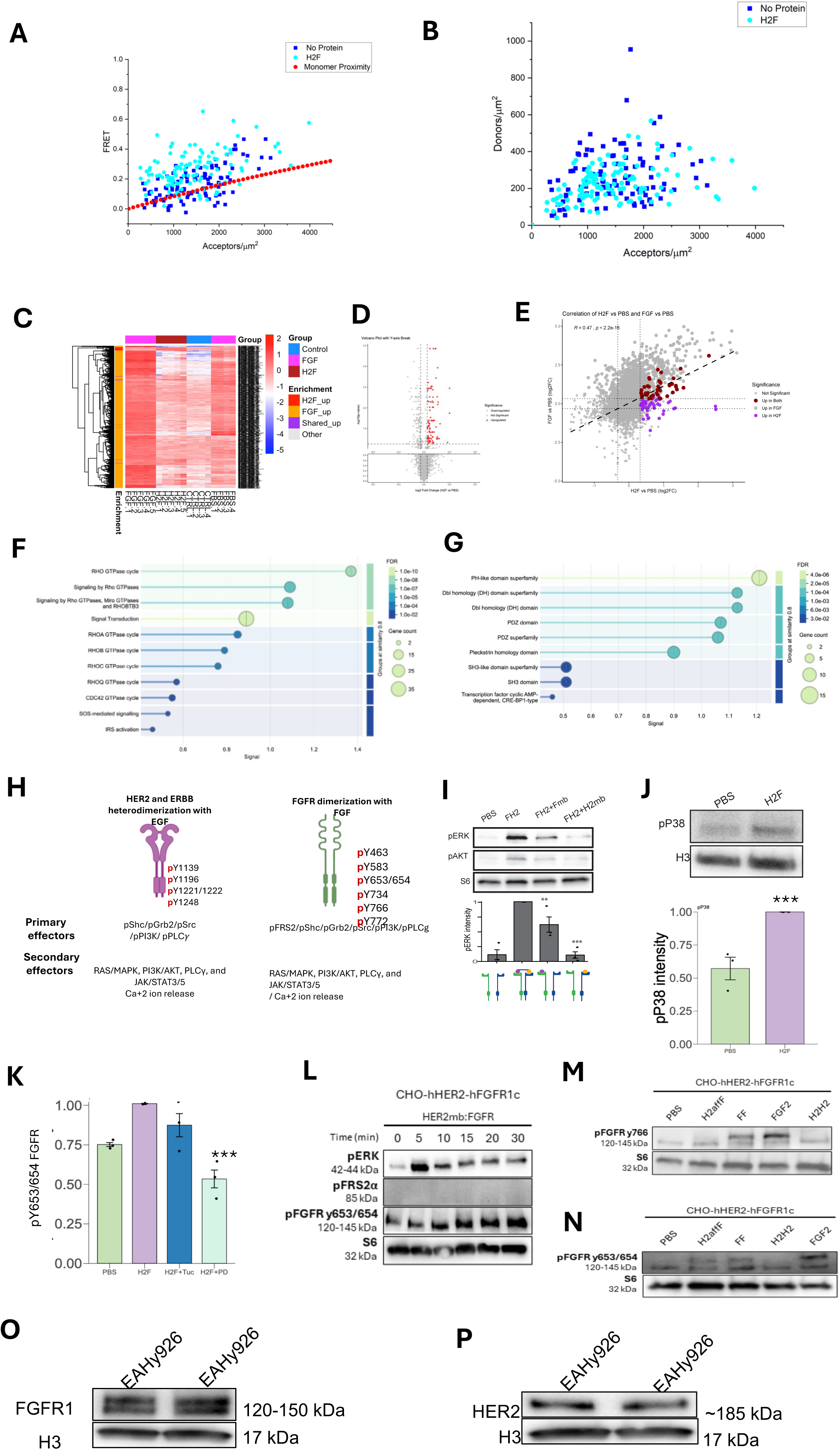
**HER2mb: FGFR1cmb requires both HER2 and FGFR1C in proximity for downstream effectors activation.** (A) FRET efficiencies and concentrations, measured in 52 cells in the absence of ligand and 61 cells in the presence of 500 nM H2F. One or two small (∼ 5 mm) regions of the plasma membrane were selected per cell, yielding 225 membrane regions total. In each region, the FRET efficiency, the donor concentration (HER-YFP), and the acceptor concentration (FGFR1c-mCherry) were measured. The FRET efficiency is plotted as a function of the acceptor concentration in (A). The red line is the “proximity FRET”, which depends only on the acceptor concentration, and accounts for random close approach of donors and acceptors in the absence of specific interactions. [37]. Single-cell FRET data lay above the red line, indicative of specific interactions. (B) The donor concentration in the FRET experiments is plotted as a function of the acceptor concentration. (C) LC/MS analysis of 6,325 unique phosphopeptides in H2F, FGF2, FBS treated CHO-hHER2-hFGFR1c cells. Heatmap represents the enrichment of phosphopeptides in each of the treatment replicates. (D) A volcano map showing the significantly enriched phosphopeptides(red dots) detected in 100nM H2F treated CHO-hHER2-hFGFR1c cells for 15 min compared to PBS control treated. (E) A correlation map shows a weak correlation between significantly H2F enriched and FGF2 enriched phosphopeptides over PBS treated. A set of phosphopeptides are overlapping (red dots) between H2F as well as FGF2 enriched while few phosphopeptides are specific to H2F (purple dots). (F) Shared phosphopeptides between H2F and FGF show GO molecular functions such as PI3K/AKT, IRS activation, PIP3 mediated AKT activation. (G) Shared phosphopeptides between H2F and FGF show intracellular signalling by second messengers. (H) Schematic showing various known natural primary and secondary downstream effectors of HER2 heterodimerization with ERBB1 in presence of natural ligand EGF (left), and FGFR dimerization in presence of natural ligand FGF (right). (I) A western blot showing H2affF shows pERK response in CHO-hHER2-hFGFR1c cells but not in CHO-hHER2 or CHO-hFGFR1c. (J) pP38 phosphorylation western blot shows enrichment in H2F treated CHO-hHER2-hFGFR1c cells. The bar graph shows the western blot quantification of pP38 in H2F treated CHO-hHER2-hFGFR1c cells. (K) Western blot showing pERK, pFRS2, pY653/654 FGFR, and S6 in HER2mb:FGFRmb(H2F) treatment overtime at various time points in CHO-hHER2-hFGFR1c cells. (L) pY653/654 residue on FGFR western blot quantification shows loss of pY653/654 upon FGFR selective kinase inhibitor PD 173074 but shows no change in tucatinib treatment which is HER2 specific inhibitor. (M) H2affF treatment on CHO-hHER2-hFGFR1c cells shows pY653/654 on FGFR (N) H2affF treatment on CHO-hHER2-hFGFR1c cells does not show spY766 phosphorylation on FGFR. (O) Western blot showing FGFR1 is expressed in EAHy926 cells. (P) Western blot showing HER2 is expressed in EAHy926 cells.

**Supplemental Figure 5.**
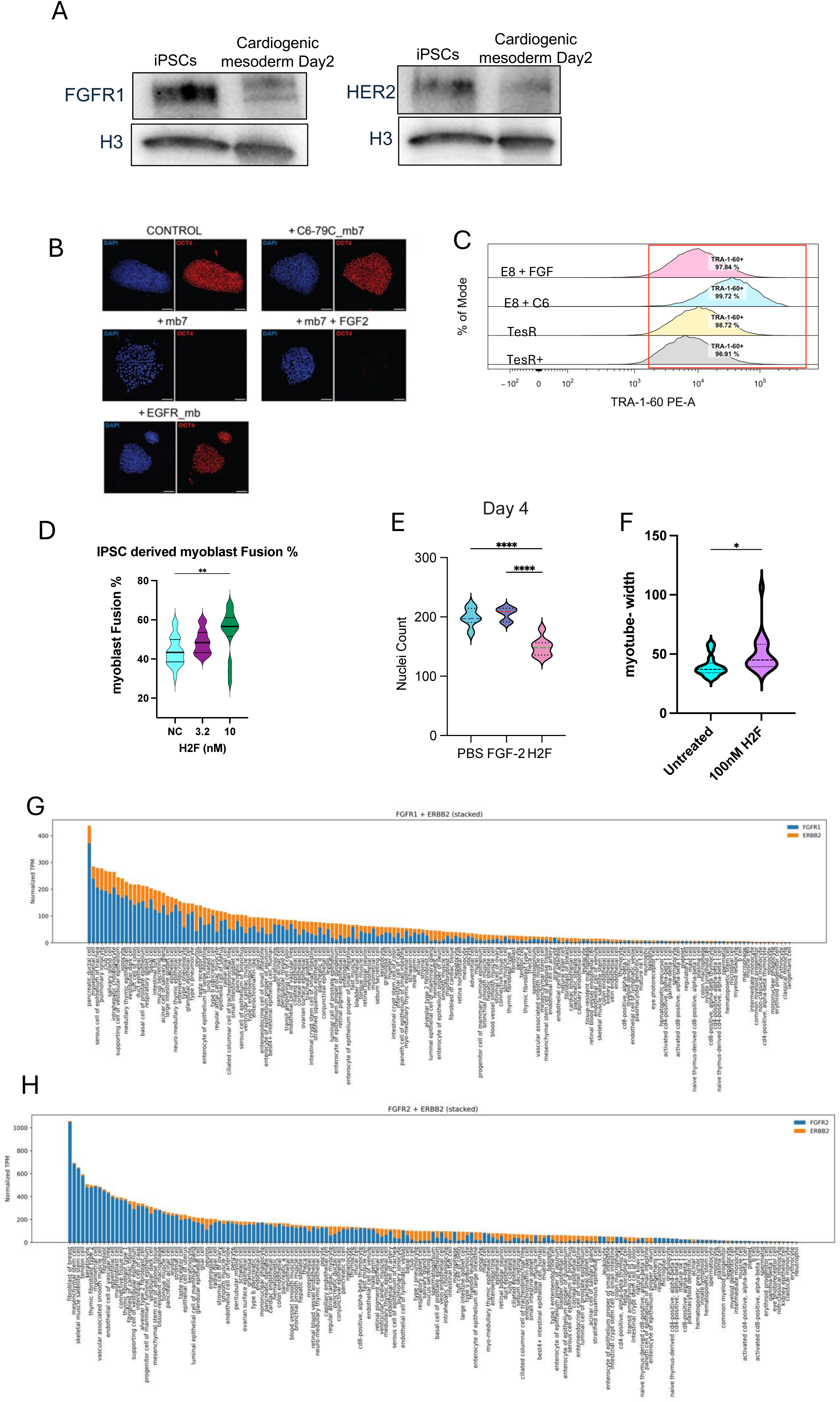
A. FGFR1 and HER2 receptors are expressed in human iPSCs as well as cardiogenic mesoderm cells (day2) where H2F is being treated in our experiments. B. Confocal images showing staining of OCT4 (red) and DAPI (blue) on mb7(FGFR1/2c antagonist), C6-79C_mb7 (FGFR1/2c agonist), EGFRmb, and FGF2 + mb7 treated iPSCs versus untreated iPSCs in mTeSR media. C. Flow cytometry data showing that C6 in minimal E8 media maintains the TRA-1-60 expression in iPSCs after 48 hours better than that of E8 + FGF, mTeSR, and mTesR+ D. Myoblast fusion % of iPSCs derived myoblast in untreated and various concentrations of H2F. E. total nuclei count in primary myoblast differentiation experiment increases in FGF treatment suggesting myoblast are proliferating under FGF growth factor treatment at da4. In H2F treatment nuclei count does not increase at day 4 suggesting H2F has a distinct mechanism and signalling than FGF that promotes differentiation of myotubes instead of proliferation. F. Primary myotube width at day 6 in 100nM H2F treatment is relatively higher than the control. G. A graph representing the normalised transcripts per million of receptors FGFR1+HER2 H. FGFR2+HER2 coexpressing cell types, single cell data analysed from Tabulea sapiens. These cell types are potential targets of H2F.

